# Genetics of human plasma lipidome: Understanding lipid metabolism and its link to diseases beyond traditional lipids

**DOI:** 10.1101/457960

**Authors:** Rubina Tabassum, Joel T. Rämö, Pietari Ripatti, Jukka T. Koskela, Mitja Kurki, Juha Karjalainen, Shabbeer Hassan, Javier Nunez-Fontarnau, Tuomo T.J. Kiiskinen, Sanni Söderlund, Niina Matikainen, Mathias J. Gerl, Michal A. Surma, Christian Klose, Nathan O. Stitziel, Hannele Laivuori, Aki S. Havulinna, Susan K. Service, Veikko Salomaa, Matti Pirinen, Matti Jauhiainen, Mark J. Daly, Nelson B. Freimer, Aarno Palotie, Marja-Riitta Taskinen, Kai Simons, Samuli Ripatti

**Author notes:** Corresponding author: Prof. Samuli Ripatti Institute for Molecular Medicine Finland (FIMM), University of Helsinki, PO Box 20, FI-00014 University of Helsinki, Finland, E-mail: samuli.

## Abstract

**Aim:** Genetic investigation of human plasma lipidome to get insights into lipid-related disorders beyond traditional lipid measures.

**Methods and Results:** We performed a genome-wide association study (GWAS) of 141 lipid species (n=2,181 individuals), followed by phenome-wide scans (PheWAS) with 44 clinical endpoints related to cardiometabolic, psychiatric and gastrointestinal disorders (n=456,941 individuals). SNP-based heritability for lipid species ranged from 0.10-0.54. Lipids with long-chain polyunsaturated fatty acids showed higher heritability and genetic sharing, suggesting considerable genetic regulation at acyl chains levels. We identified 35 genomic regions associated with at least one lipid species (P<5×10^−8^), revealing 37 new SNP-lipid species pair associations e.g. new association between *ABCG5/8* and CE(20:2;0). PheWAS of lipid-species-associated loci suggested new associations of *BLK* with obesity, *FADS2* with thrombophlebitis, and *BLK* and *SPTLC3* with gallbladder disease (false discovery rate <0.05). The association patterns of lipid-species-associated loci supplied clues to their probable roles in lipid metabolism e.g. suggestive role of *SYNGR1, MIR100HG*, and *PTPRN2* in desaturation and/or elongation of fatty acids. At known lipid loci (*FADS2, APOA5* and *LPL*), genetic associations provided detailed insights to their roles in lipid biology and diseases. We also show that traditional lipid measures may fail to capture lipids such as lysophospatidylcholines (LPCs) and phosphatidylcholines (PCs) that are potential disease risk factors, but are not included in routine screens. The full genome-wide association statistics are available on the web-based database (http://35.205.141.92).

**Conclusion:** Our study reveals genetic regulation of plasma lipidome and highlights the potential of lipidomic profiling in disease gene mapping.

## Introduction

Plasma lipids are risk factors for various complex disorders like cardiovascular disease (CVD) and type 2 diabetes.^1,2^ Low-density lipoprotein cholesterol (LDL-C), high-density lipoprotein cholesterol (HDL-C), non-HDL-C, lipoprotein (a), total triglycerides and total cholesterol are “traditional lipid measures” routinely used to dissect dyslipidemia and risks for diseases. Human plasma comprises of hundreds of chemically and functionally diverse molecular lipid species,^3^ which are not captured in everyday, clinical lipid analysis. Lipid species belonging to cholesterol esters (CE), lysophosphatidylcholines (LPC), phosphatidylcholines (PC), phosphatidylethanolamines (PE), ceramides (CER), sphingomyelins (SM), and triacylglycerols (TAG) species have been shown to improve risk assessment over traditional lipid measures and other risk factors for type 2 diabetes and CVD events.^4–9^ Moreover, the fatty acid compositions of phospholipids have been implicated in coronary heart disease beyond traditional lipid measures.^10^

Genetic screens have identified over 250 genomic loci associated with traditional lipid levels.^11,12^ These genetic findings have helped to understand lipid biology and physiological processes underlying CVD and other diseases. For the majority of the genomic loci, however, the causal genes and/or their effects on detailed lipidomes beyond traditional lipid measures are unknown. Only a few studies have reported genetic associations for lipid species either through studies on subsets of the lipidome ^13, 14^ or GWASs on metabolomic measures.^15–20^ In most of these studies, however, the lipids have not been resolved on the molecular level of fatty acid or acyl chain composition (molecular lipid species).

We considered that the genetic investigation of detailed lipidomic profiles could provide better insight into lipid metabolism and its link to clinical outcomes surpassing traditional lipid measures. To test this, we carried out a GWAS of lipidomic profiles of 2,181 individuals followed by PheWAS with 44 clinical end-points related to cardiometabolic, psychiatric and gastro-intestinal disorders in an independent dataset of 456,941 individuals from the Finnish and UK Biobanks (Figure 1). We aimed to identify genomic loci influencing plasma levels of lipid species and their effects on the risk of diseases. We also set out to answer the following questions: (1) how heritable are the lipid species and do they share genetic components (2) can we gain mechanistic insights to pathways linking genetic variation to disease risk through detailed measures of the lipid species and (3) could detailed lipid profiles provide additional biological insights to genetic regulation of lipid metabolism for the previously identified lipid loci.

**Figure 1:**
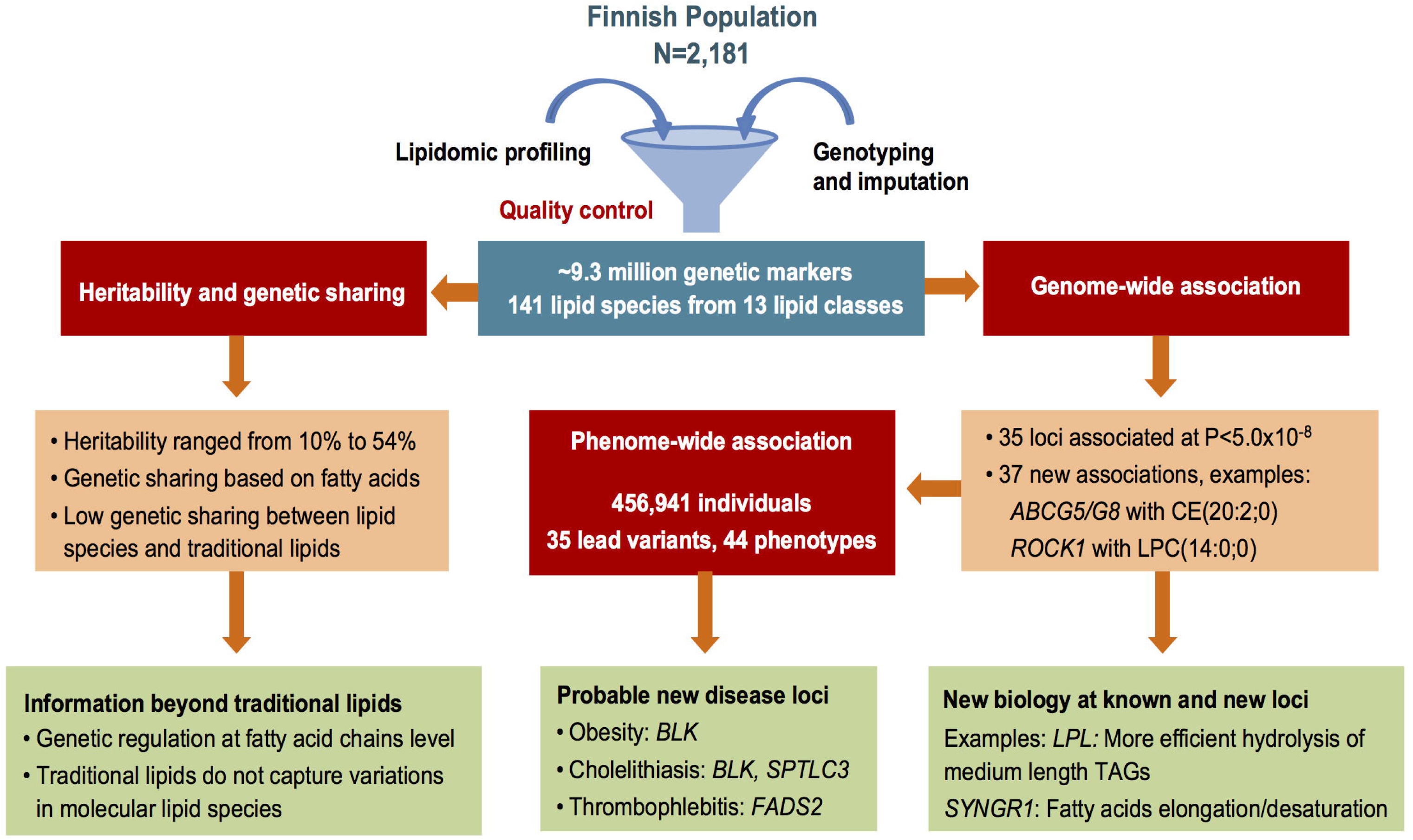
Genetics of human plasma lipidome. The figure illustrates the study design and key findings of the study.

## Methods

### Study cohorts

The detailed description of the subject recruitment and measurements is provided in Supplementary Data. Briefly, the study included participant from the following cohorts:

#### EUFAM

The European Multicenter Study on Familial Dyslipidemias in Patients with Premature Coronary Heart Disease (EUFAM) study cohort is comprised of the Finnish familial combined hyperlipidemia families.^21^ The families in EUFAM study were identified via probands admitted to Finnish university hospitals with a diagnosis of premature coronary heart disease. For the lipidomic profiling, 1,039 EUFAM participants for which serum samples were available were included (Supplementary Table 1).

#### FINRISK

The Finnish National FINRISK study is a population-based survey conducted every 5 years since 1972 (detailed in Supplementary Data).^22^ Lipidomic profiling was performed for 1,142 participants that were randomly selected from the FINRISK 2012 survey (Supplementary Table 1).

#### Finnish Biobank

The Finnish Biobank data is composed of 47,980 Finnish participants with 807 phenotypes derived from ICD codes in Finnish national hospital registries and cause-of-death registry as a part of FinnGen project.

#### UK Biobank

The UK Biobank data is comprised of >500,000 participants based in UK and aged 40–69 years, annotated for over 2,000 phenotypes.^23^ The PheWAS analyses in the present study included 408,961 samples from white British participants.

Written informed consent was obtained from all the study participants. The study protocols were approved by the ethics committees of the participating centers. The study was conducted in accordance with the principles of the Helsinki declaration.

### Lipidomic profiling

Mass spectrometry-based lipid analysis of 2,181 participants was performed in three batches-353 and 686 EUFAM participants in two batches and 1,142 FINRISK participants in third batch at Lipotype GmbH (Dresden, Germany) as detailed in Supplementary Data. Data were corrected for batch and drift effects. Lipid species detected in <80% of the samples in any of the batches and samples (N=64) with low lipid contents were excluded. A total of 141 lipid species from 13 lipid classes (Supplementary Table 2) detected consistently in three batches were included in further analyses. The total amounts of lipid classes were calculated by summing up respective lipid species. The measured concentrations of the lipid species and calculated class total were transformed to normal distribution by rank-based inverse normal transformation.

### Genotyping and imputation

Genotyping for both EUFAM and FINRISK cohorts was performed using the HumanCoreExome BeadChip (Illumina Inc., San Diego, CA, USA). Genotype data underwent stringent quality control (QC) before imputation using an in-house QC pipeline (Supplementary Data). Imputation was performed using IMPUTE2,^24^ which used two population specific reference panels of 2,690 high coverage whole-genome and 5,093 high coverage whole exome sequence data. Variants with imputation info score <0.70 were filtered out. After QC on lipidomic profiles and imputed variants, all subsequent analyses included 2045 individuals and ~9.3 million variants with MAF >0.005 that were available in both cohorts.

Genotyping of the Finnish Biobank samples was performed using arrays from Illumina and Affymetrix (Santa Clara, CA, USA) and custom-designed Affymetrix array for the FinnGen project. Quality control and imputation were performed using the same pipeline as described above. Genotyping for the majority of the UK Biobank participants was done using the Affymetrix UK Biobank Axiom Array, while a subset of participants was genotyped using the Affymetrix UK BiLEVE Axiom Array. Details about the quality control and imputation of UK Biobank cohort are described earlier.^25^

### Heritability estimates and genetic correlations

For heritability and genetic correlation estimation, rank-based inverse transformed measures of lipid species, computed separately for the EUFAM and FINRISK cohorts, were combined to increase statistical power. The residuals of transformed measures after regressing for age, sex, first ten principal components (PCs) of genetic population structure, lipid medication, hormone replacement therapy, thyroid condition and type 2 diabetes were used as phenotypes. A genetic relationship matrix (GRM) was generated using GCTA,^26^ after removing variants with missingness >10% and MAF <0.005. The generated GRM was then used to estimate heritability using variance component analysis and genetic correlations using bivariate linear mixed model as implemented in biMM.^27^ The heritability estimates of lipid species in different groups were compared using Mann-Whitney *U* test. The phenotypic correlations based on the plasma levels between all the pairs of the lipid species and traditional lipid measures were calculated using Pearson’s correlation coefficient. The heatmaps and hierarchical clustering based on genetic and phenotypic correlations were generated using heatmap.2 in R.

### Lipidomics GWAS and meta-analysis

We performed univariate association tests for 141 individual lipid species, 12 total lipid classes and 4 traditional lipid measures (HDL-C, LDL-C, total cholesterol and triglycerides), in all batches to control for possible batch effects and combined the summary statistics by meta-analysis. The association analyses for the EUFAM cohort were performed using linear mixed models including the above-mentioned covariates as fixed effects and kinship matrix as random effect as implemented in MMM.^28^ The kinship matrices for the GWAS analyses were computed separately for each chromosome to include the variants from the other chromosomes using directly genotyped variants with MAF >0.01 and missingness <2%. The FINRISK cohort was analyzed with linear regression model adjusting for age, sex, first ten PCs, lipid medication and diabetes using SNPTEST v2.5.^29^ Meta-analyses were performed using the inverse variance weighted method for fixed effects adjusted for genomic inflation factor in METAL.^30^ In addition, analyses adjusting for the traditional lipid measures (in addition to above-mentioned covariates) were also performed for the identified variants to determine the independent effect on lipid species.

Test statistics were adjusted for λ values if >1.0 before meta-analyses. Genomic inflation factor (λ) ranged from 0.98-1.19 across the batches whereas the final λ values for meta-analysis ranged from 0.998 to 1.045 (Supplementary Table 3). Associations with P value <5.0×10^−8^ and consistent in direction of effect in all three batches were considered significant. Variants were designated as novel if not located within 1Mb of any previously reported variants for lipids (any of the traditional lipid measures and molecular lipid species); and as independent signal in known locus if located within 1 Mb but r^2^<0.20 with the previous lead variants and confirmed by conditional analysis. Variants with the strongest association in the identified lipid species loci was identified as the lead variants and were annotated to the nearest gene for the new loci.

### PheWAS

We identified 95 disease phenotypes that have previously been linked to lipid levels including cardiometabolic, psychiatric and gastrointestinal disorders from the derived phenotypes in the Finnish Biobank. We manually mapped 44 of these 95 phenotypes to UK Biobank phenotypes (Supplementary Table 4). For the Finnish cohort, the associations between the 35 lead variants from the identified loci and these 44 phenotypes were obtained from the ongoing PheWAS as a part of the FinnGen project. The associations were tested using saddle point approximation method adjusting for age, sex, and first 10 PCs as implemented in SPAtest R package.^31^ The association between these 44 phenotypes and 35 lead variants in UK Biobank were obtained from Zhou et al. that were tested using logistic mixed model in SAIGE with a saddle point approximation and adjusting for first four principal components, age and sex (https://www.leelabsg.org/resources).^32^ In addition, associations with obesity and body mass index (BMI) were also tested using logistic and linear regression models respectively with the same covariates as mentioned above, both for Finnish and UK Biobank cohorts. Individuals with BMI ≥30kg/m^2^ were categorized as obese. Metaanalyses of both cohorts were performed using the inverse variance weighted method for fixed effects model in METAL. All the PheWAS associations with false discovery rate (FDR) <5% evaluated using the Benjamini-Hochberg method were considered significant.

### Variance explained

To determine the variance explained by the known loci for traditional lipid measures, we included all the lead variants with MAF >0.005 in 250 genomic loci that have previously been associated with one or more of the four traditional lipid measures (Supplementary Table 5). Of the 636 reported variants, 557 variants with MAF >0.005 (including six proxies) were available in our QC passed imputed genotype data. A genetic relationship matrix (GRM) based on these 557 variants was generated using GCTA that was used to determine the variance in plasma levels of all lipid species explained by the known variants using variance component analysis in biMM.

## Results

### Genetic contribution to lipidome

SNP based heritability estimates ranged from 0.10 to 0.54 (Figure 2A, Supplementary Table 2), with considerable variation across lipid classes (Figure 2B), with similar trends as reported previously.^33,34^ Ceramides (CER) showed the greatest estimated heritability (median=0.30, IQR=0.09) whereas phosphatidylinositols (PIs) showed the least heritability (median=0.19, IQR=0.045). Sphingolipids had higher heritability than glycerolipids ranging from 0.27 to 0.41 (Figure 2B), which is similar to a previous study that reported higher heritability for sphingolipids ranging from 0.28 to 0.53 estimated based on pedigrees.^33^

**Figure 2:**
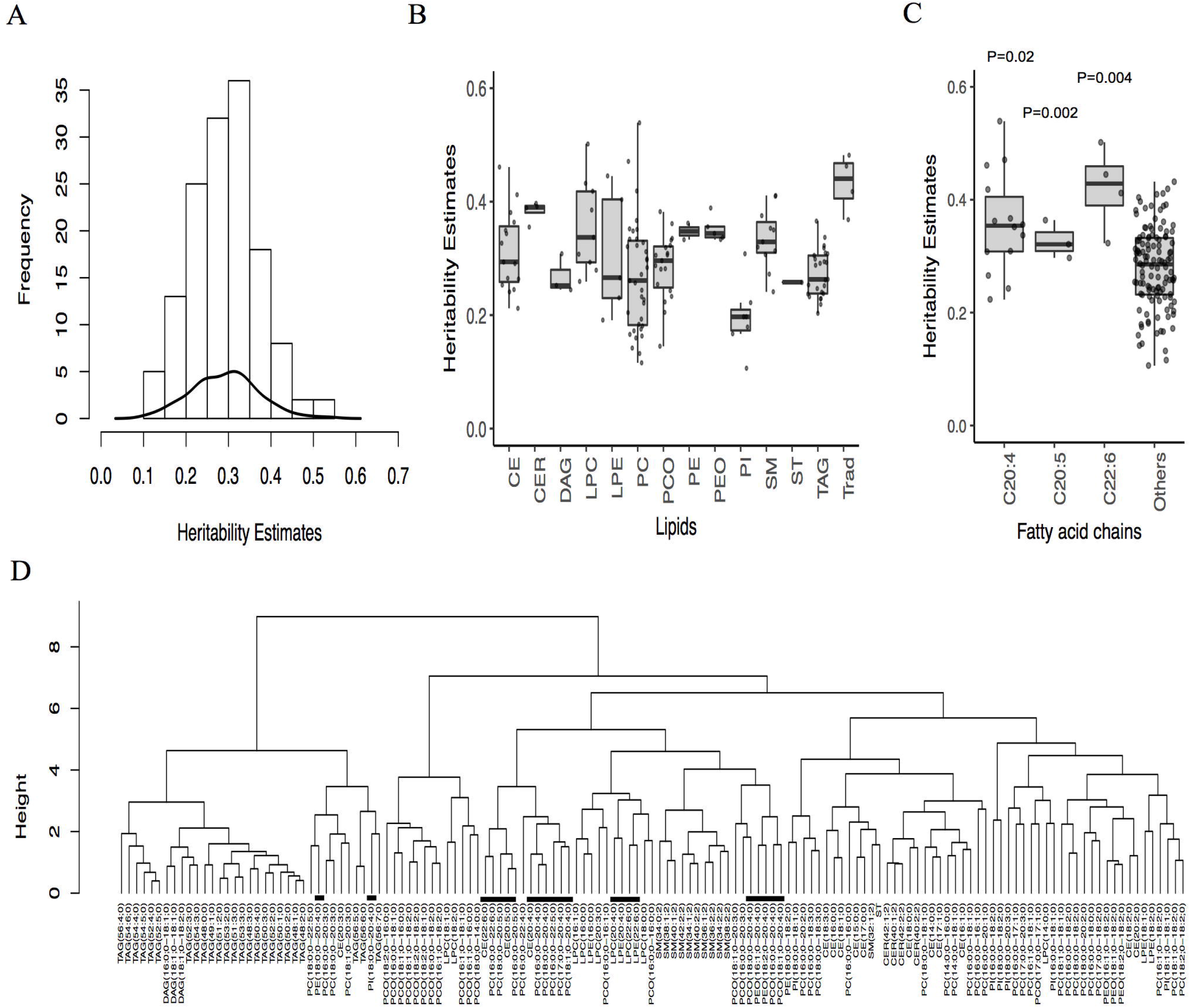
Heritability of lipidomic profiles and genetic correlations among the lipid species. (A) Histogram and kernel density curve showing the distribution of heritability estimates across all the lipid species. (B) Boxplot showing the medians and ranges of heritability estimates in each lipid class. (C) Boxplot showing comparison of the median heritability estimates of lipid species containing C20:4, C20:5 and C22:6 acyl chains and all others. (D) Hierarchical clustering of lipid species based on genetic correlations showing genetic sharing among lipid species. Lipids containing polyunsaturated fatty acids C20:5, C20:4, and C22:6 are highlighted with black bars. CER=Ceramide, DAG=Diacylglyceride, LPC=Lysophosphatidylcholine, LPE=Lysophosphatidylethanolamine, PC=Phosphatidylcholine, PCO=Phosphatidylcholine-ether, PE=Phosphatidylethanolamine, PEO=Phosphatidylethanolamine-ether, PI=Phosphatidylinositol, CE=Cholesteryl ester, SM=Sphingomyelin, ST=Sterol, TAG=Triacyglycerol.

Lipids containing polyunsaturated fatty acids, particularly C20:4, C20:5 and C22:6, had significantly higher heritability compared to other lipid species (Figure 2C). For instance, PC(17:0;0-20:4;0) and LPC(22:6;0) had the highest heritability (>0.50) whereas PC(16:0;0-16:1;0) and PI(16:0;0-18:2;0) had the lowest heritability estimates (<0.12) (Supplementary Table 2). Moreover, longer, polyunsaturated lipids (those with four or more double bonds) had strong genetic correlations with each other than with other lipid species (Supplementary Figure 1, Supplementary Table 6). This can be seen in the hierarchical clustering based on genetic correlations that segregate TAG subspecies into two clusters based on carbon content and degree of unsaturation (Figure 2D). These patterns were not seen in phenotypic correlations that were estimated based on the plasma levels of lipid species (Supplementary Figure 2).

We observed low phenotypic and genetic correlation between traditional lipids and molecular lipid species, except strong positive genetic correlations of triglycerides with TAG and DAG subspecies (average r=0.88) (Figure 3). However, triglycerides had low genetic correlation with other lipid species (average (abs) r= 0.26). HDL-C and LDL-C levels had low genetic and phenotypic correlations with most of the lipid species (Figure 3, Supplementary Table 6). This implies that genetic variants associated with traditional lipid measures would fail to capture the variations in plasma levels of many lipid subspecies. Consistently, all of the known lipid variants explained 2% to 21% of variances in plasma levels of various lipid species, with the least variance accounting for LPCs (Figure 3).

**Figure 3:**
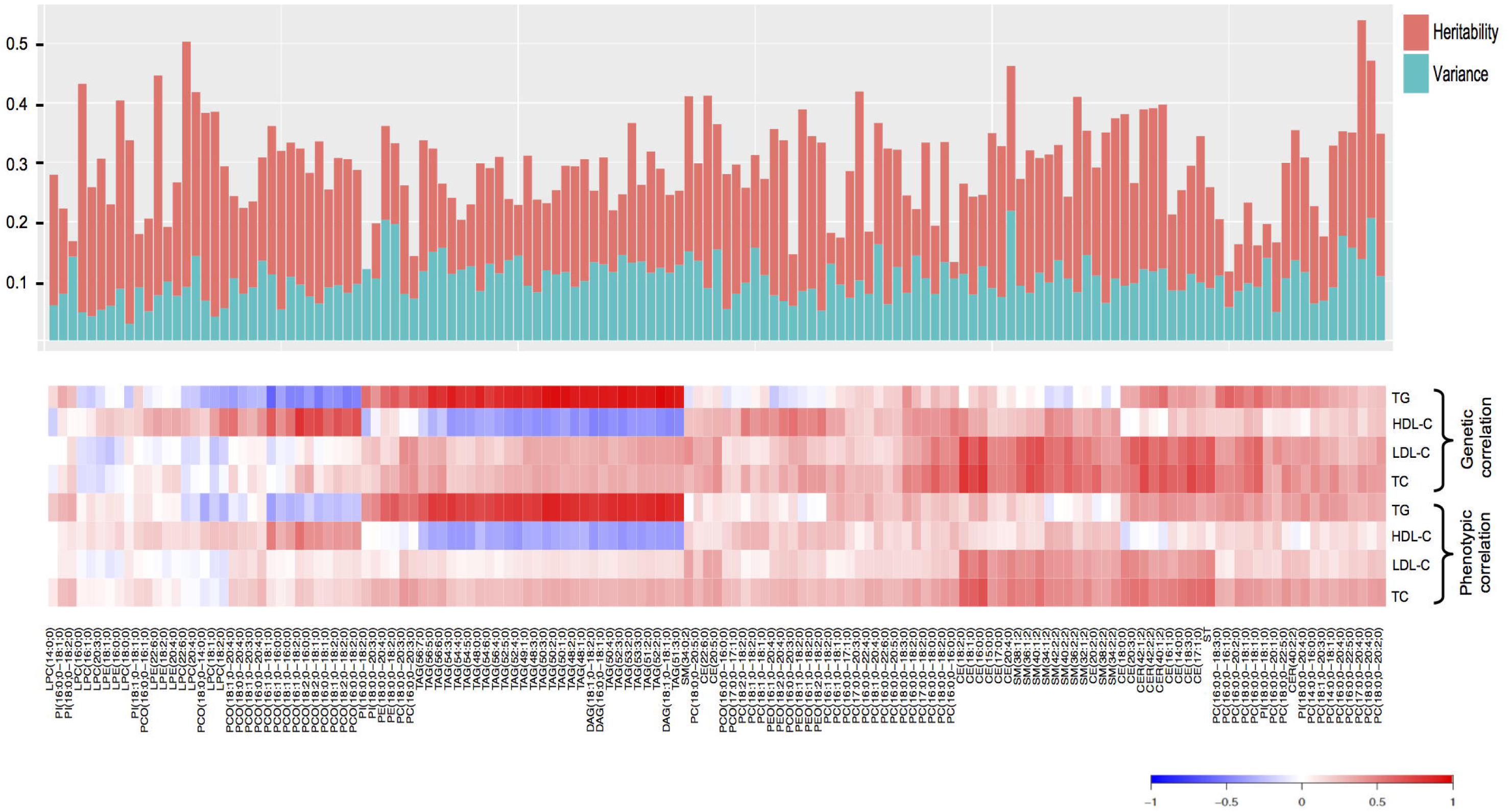
Lipidomic profiles capture information beyond traditional lipid measures. The genetic and phenotypic correlations between traditional lipid measures and molecular lipid species are shown in lower panel. The bar plot in the upper panel shows the heritability estimates of each lipid species (red bars) and the variance explained by all the known loci together (green bars). The lipid species are ordered as in lower panel.

It is to be noted that this sample size might not provide sufficient power for heritability estimations in unrelated samples. However, our study also included the family samples which provides higher statistical power in heritability estimation than unrelated samples. Moreover, lipid species with genome-wide significant association had higher heritability estimates compared to the lipid species with no significant association (Supplementary Figure 3).

### Genetic architecture of lipidome

We found 3,754 associations between 820 variants located within 35 genomic loci (1MB blocks) and 74 lipid species from 11 lipid classes at genome-wide significance level (P<5.0×10^−8^) (Supplementary Table 7). These included 37 new locus-lipid species pair associations (Figure 4). The identified loci included 3 loci for CEs, 6 loci for LPCs, 5 loci for LPEs, 4 loci for PCs, 3 for PC-Os, 2 loci for PEs, 3 loci for PE-O, 4 loci for PIs, 4 loci for SMs, 10 for TAGs and 2 loci for ceramides (Table 1, Figure 4). In line with our observation of higher heritability for lipids with C20:4, C20:5 or C22:6 acyl chains, we detected associations for 18 out of 21 lipids with these acyl chains. Moreover, 11 out of 35 identified loci were associated with at least one of the polyunsaturated lipids. The genome-wide association statistics for all lipid species are available on the web-based database (http://35.205.141.92) for mining and visualization.

**Figure 4:**
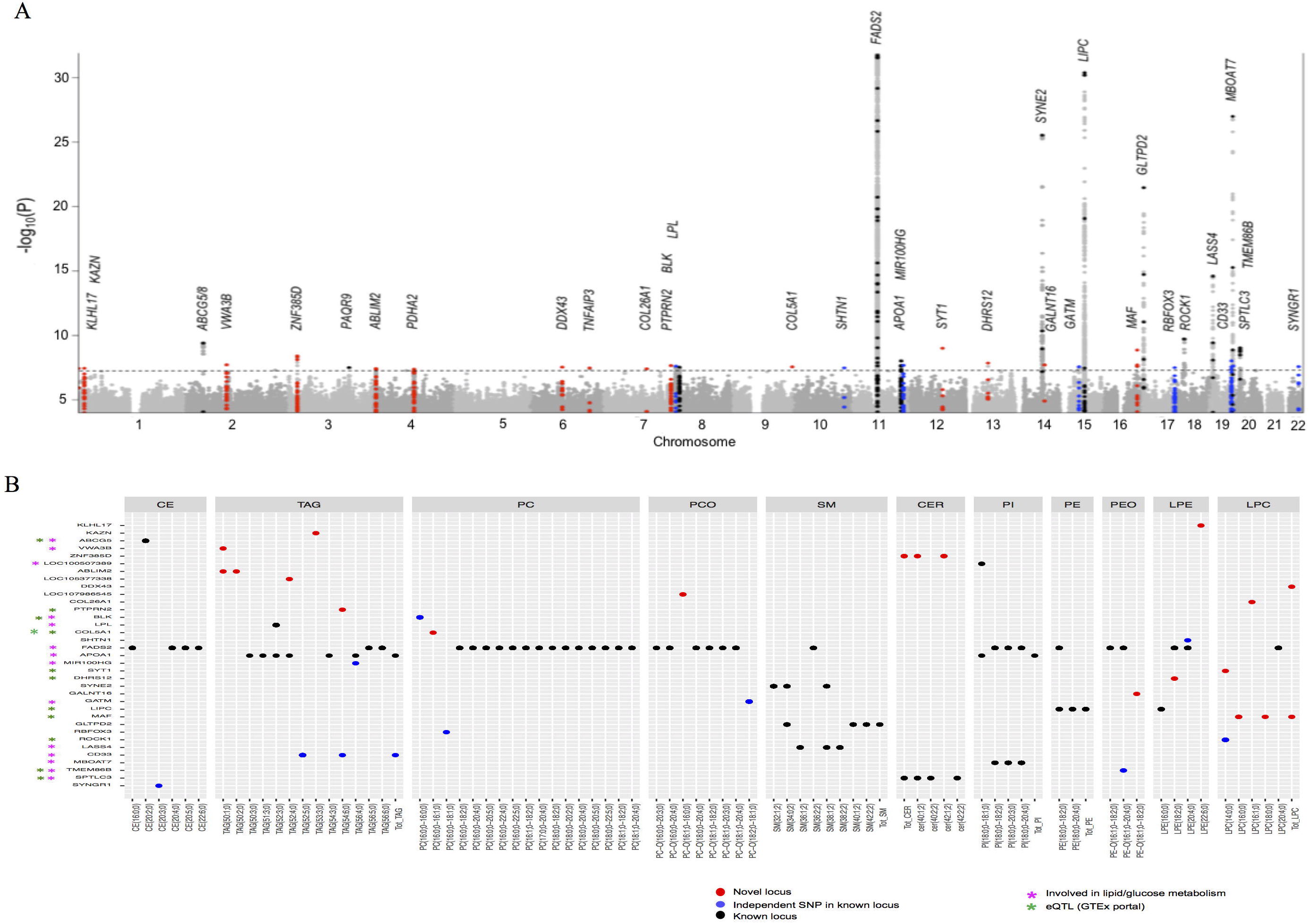
Genetic architecture of lipidome. (A) Manhattan plot showing associations for all 141 lipid species. Only the associations with P<1.0×10^−4^ and consistent directions in all three batches are plotted. Y-axis is truncated for better representation of the data. The dotted line represents the threshold for genome-wide significant associations at P<5.0×10^−8^. (B) Genome-wide significant associations between the identified lipid species associated loci and lipid species showing effect of the loci on lipidome. The associations presented with blue dots are novel whereas previously reported associations are presented in red dots. Novel hits with P<5.0×10^−8^ are shown as red dots, new independent hits in previously reported loci are presented as blue dots and hits in previously known loci are presented as black dots.

**Table 1:**
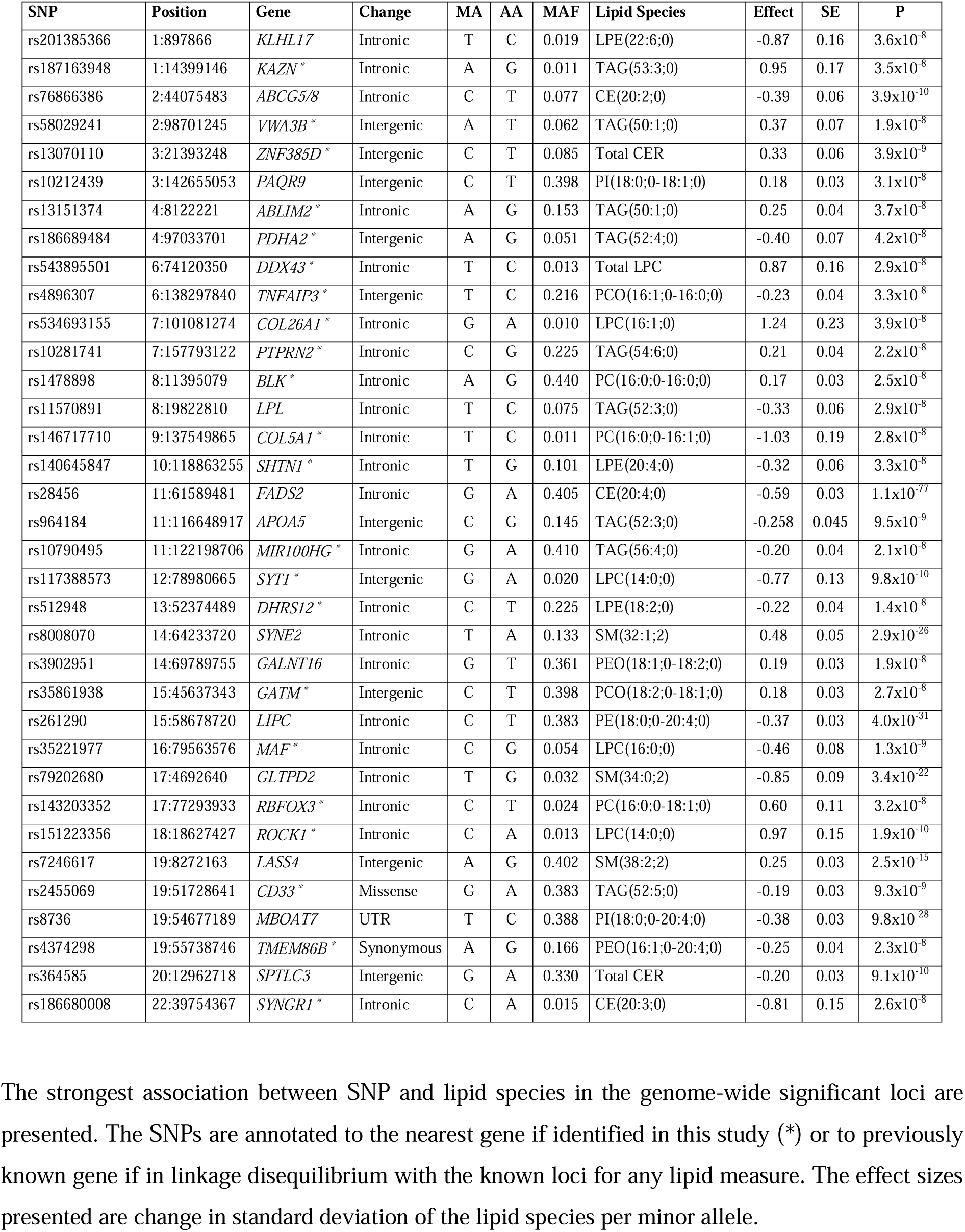
Genomic loci with molecular lipid species at genome-wide significance.

Among the 35 identified loci, 15 loci were located in genomic regions not previously reported for any lipid measure or related metabolite, and 9 loci were located near known loci for lipids but independent of any previously reported variant (Supplementary Table 8). Of these 24 new loci, 8 loci (*ROCK1*, *PTPRN2*, *MAF*, *COL5A1*, *SYT1*, *DHRS12, TMEM86B* and *BLK*) were located in or near genes that are biologically related to lipid/glucose metabolic pathways suggesting biological significance. The lead variants with the most significant SNP-lipid species associations at the identified loci are presented in Supplementary Table 8, with their genotype-phenotype relationships in Supplementary Figure 4. The strongest new association was at an intronic variant rs151223356 near *ROCK1* with short acyl-chain LPC(14:0,0) (P=1.9×10^−10^). The SNP is located near *ABHD3* locus which has previously been shown to associate with a phosphatidylcholine species PC(32:2;0),^18^ but the identified variant is independent of all previously reported variants within 1 Mb of the region (r2<0.01 and P_condtional_=2.3×10^−10^). *ROCK1* encodes for a serine/threonine kinase that plays key role in glucose metabolism and has previously been implicated in CVD and diabetes pathogenesis.^35,36^

We also identified new locus-lipid species pair associations at previously reported lipid loci, revealing their specific associations. For example, at *ABCG5/8*, a previously known loci for total cholesterol, LDL-C and CE in LDL,^11,16^ we found a novel association of *ABCG5/8* with CE(20:2;0) (P=3.9×10^−10^). At *LPL*, previously associated with plasma TG levels,^11^ our analysis revealed the strongest association of *LPL* variant with TAG(52:3;0). Similarly, at *MBOAT7* and *LIPC*, we identified new and specific associations with PI(18:0;0-20:4;0) and PE(18:0;0-20:4;0) respectively. Moreover, we replicated previous associations of *FADS2*, *SYNE2*, *LIPC*, *GLTPD2*, *LASS4* and *MBOAT7*,^13–20^ either with the same lipid species or the same lipid class.

### Lipid species associated loci and metabolic pathways

Based on the patterns in the associations of the identified loci with various lipid species (Supplementary Figure 5), we propose mechanisms for their effects on lipid metabolism. With a few examples, we show how lipidomic profiles could help to further increase our understanding of lipid biology and gene functions as detailed below. The *FADS2* rs28456-G was associated with increased levels of lipids with C20:3 acyl chain and decreased levels of lipids with C20:4, C20:5, and C22:6 acyl chains (Supplementary Figure 5). This inverse relationship in lipids with different polyunsaturated fatty acids (PUFAs) could be explained by the inverse effect of rs28456-G on *FADS1* and *FADS2* expression [GTEx v7]. The rs28456-G increases *FADS2* expression that might result in increased desaturation of linoleic acid (C18:2, n-6) and alpha-linolenic acid (C18:3, n-3) and hence increased dihomo-gamma-linolenic acid (C20:3, n-6) (Figure 5). Interestingly, *FADS2* rs28456-G decreases *FADS1* expression that might reduce delta 5 desaturation and subsequently resulting in decreased arachidonic acid (C20:4, n-5), eicosapentaenoic acid (C20:5, n-3), and docosahexaenoic acid (C22:6, n-3).

Furthermore, we found that *APOA5* rs964184-C and *LPL* rs964184-T were associated with reduced levels of medium length TAGs (C50 to C56), with strongest associations with TAG(52:3;0). The striking similar patterns of associations of *APOA5* rs964184-C and *LPL* rs11570891-T with TAG species suggest that both these variants might lead to more efficient hydrolysis of medium length TAGs (Figure 5). To test this, we determined the effect of *LPL* rs11570891-T on LPL enzymatic activity and relationship between LPL activity and TAG subspecies using post-heparin LPL measured in EUFAM cohort. We found that *LPL* rs11570891-T (an eQTL increasing *LPL* expression) was associated with the increased LPL activity which in turn was associated with TAG species with higher effect on medium length TAGs than other TAGs (Figure 5).

**Figure 5:**
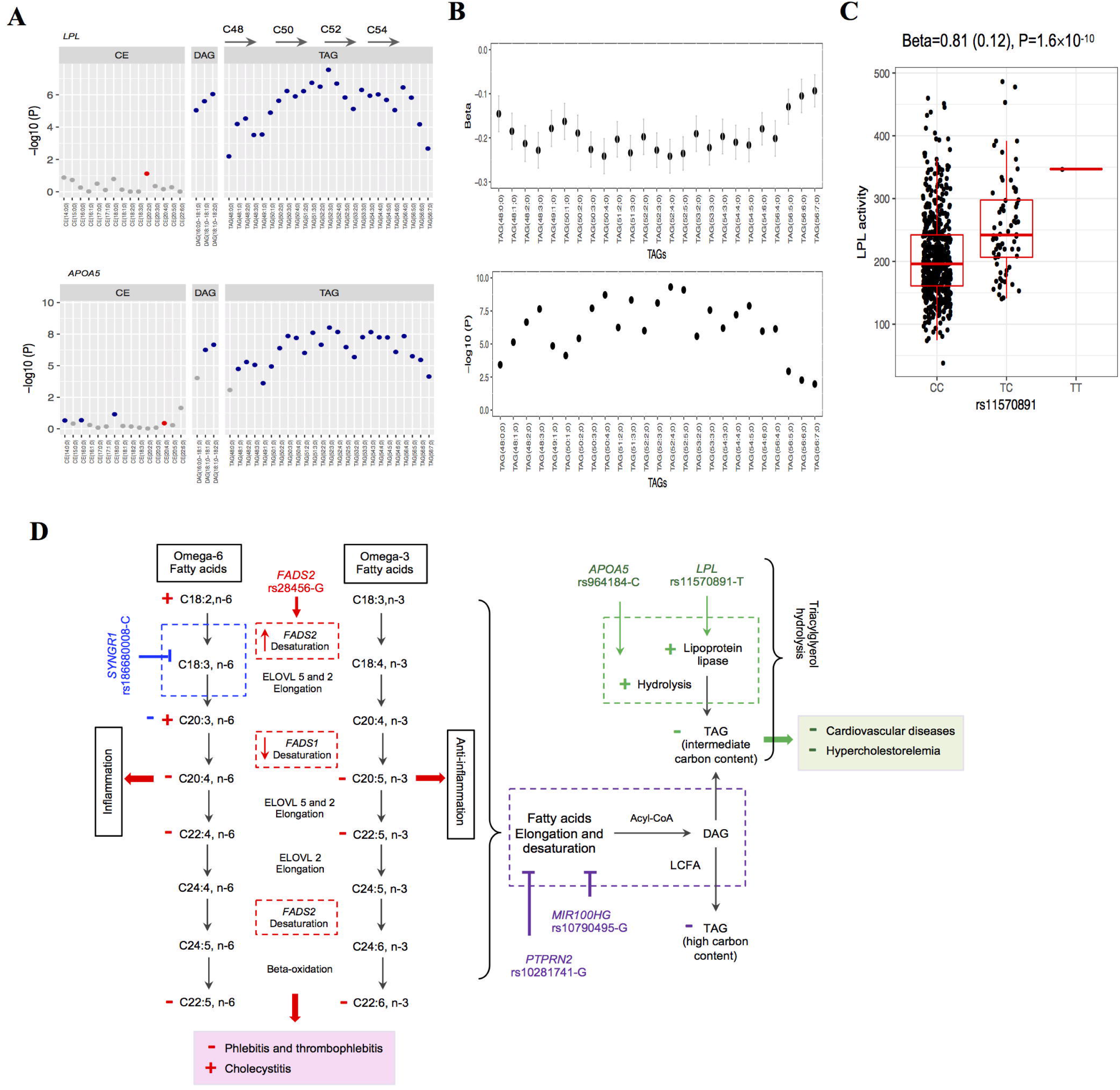
Patterns in associations and proposed mechanisms for the effect of the selected lead variants from lipid species associated loci on lipid metabolism and clinical outcomes. (A) P values for associations of *LPL* and *APOA5* variants with TAGs showing patterns in associations. The data points are color coded by the direction of effect on lipid species-increased level (red), decreased level (blue) and inconsistent direction across the cohorts (grey). (B) Association of LPL activity with TAGs. The change (Beta ±SE) in plasma levels of different TAGs with per increase in standard deviation of LPL activity (upper panel) and their respective P values (lower panel) in 978 individuals are plotted. (C) Association of *LPL* variant rs11570891 with LPL activity. Plotted are the measured LPL activity after 15mins of heparin load for 626 individuals with different genotypes. (D) Based on the patterns of the association of lipid species associated loci with different lipid species, we propose that: (1) *LPL* rs11570891-T and *APOA5* rs964184-C might result in more efficient hydrolysis of medium length TAGs which might results in reduced CVD risk (2) *FADS2* rs28456-G have observed effect on PUFA metabolism (shown by red colored + and – signs) through its inverse effect on *FADS2* and *FADS1* expressions (2) *SYNGR1* rs18680008-C might have role in negative regulation of either desaturation of linoleic acid (C18:2,n-6) or elongation of gamma linoleic acid (C18:3,n-6) which could lead to decreased levels of lipids with C20:3 (blue color). (4) *PTPRN2* rs10281741-G and *MIR100HG* rs10790495-G, which have very similar patterns of association with reduced level of long chain polyunsaturated TAG species, might have role in negative regulation of either elongation and desaturation of fatty acids or incorporation of long chain unsaturated fatty acids in glycerol backbone during TAG biosynthesis (purple color). The + and -signs indicate increase or decrease in the level of lipid species or risk of disease respectively as observed in our study, with different colors for different variant.

*SYNGR1* rs186680008-C showed stronger associations with decreased levels of lipid species with C20:3 acyl chain from different lipid classes including CEs, PCs and PCOs (Supplementary Figure 5). This suggests that *SYNGR1* rs18680008-C might have role in negative regulation of either desaturation of linoleic acid (C18:2, n-6) or elongation of gamma linoleic acid (C18:3, n-6) which could lead to decreased levels of lipids with C20:3 acyl chain, presumably dihomo-gamma-linolenic acid (DGLA; C20:3, n-6) (Figure 5). *PTPRN2* rs10281741-G and *MIR100HG* rs10790495-G showed very similar patterns of associations with reduced levels of long polyunsaturated TAG species, suggesting their role in negative regulation of either elongation and desaturation of fatty acids or incorporation of long chain unsaturated fatty acids during TAG biosynthesis.

### Lipid species associated loci and disease risks

PheWAS revealed associations of lead variants from six lipid species associated loci (*APOA5*, *ABCG5/8*, *BLK*, *LPL*, *FADS2* and *SPTLC3*) with at least one of the clinical endpoints (FDR<5%) (Table 2, Supplementary Table 9). These included novel associations of variants at *FADS2*, *BLK* and *SPTLC3* with various disease outcomes. *FADS1-2-3* is a well-known lipid modifying locus, however, like many other known lipid loci, its effects on CVD risk has been unclear. We found association of *FADS2* rs28456-G with lower risk of phlebitis and thrombophlebitis.

**Table 2:** Association of lipid species associated loci with disease outcomes

| Gene | SNP | Phenotype | Effect allele | Other allele | Effect on Phenome<sup>Ψ</sup> | P value | Effect on lipidome # | Effect on transcriptome* |
| --- | --- | --- | --- | --- | --- | --- | --- | --- |
| *ABCG5/8* | rs76866386 | Cholelithiasis | C | T | 2.06 (2.01-2.10) | 9.8x10<sup>-238</sup> | CE(20:2;0) - | *ABCG5* + (spleen) |
| *ABCG5/8* | rs76866386 | Acute pancreatitis | C | T | 1.46 (1.25-1.76) | 2.2x10<sup>-10</sup> | CE(20:2;0) - | *ABCG5* + (spleen) |
| *ABCG5/8* | rs76866386 | Hypercholesterolemia | C | T | 0.90 (0.87-0.93) | 1.4x10<sup>-9</sup> | CE(20:2;0) - | *ABCG5* + (spleen) |
| *ABCG5/8* | rs76866386 | Coronary atherosclerosis | C | T | 0.92 (0.89-0.96) | 1.1x10<sup>-4</sup> | CE(20:2;0) - | *ABCG5* + (spleen) |
| *BLK* | rs1478898 | Hypertension | A | G | 0.96 (0.95-0.98) | 2.1x10<sup>-9</sup> | PC(16:0;0-16:0;0) + | *BLK* + (fibroblasts) |
| *BLK* | rs1478898 | Obesity | A | G | 0.97 (0.96-0.98) | 5.6x10<sup>-8</sup> | PC(16:0;0-16:0;0) + | *BLK* + (fibroblasts) |
| *BLK* | rs1478898 | Cholelithiasis | A | G | 1.06 (1.03-1.08) | 1.9x10<sup>-6</sup> | PC(16:0;0-16:0;0) + | *BLK* + (fibroblasts) |
| *LPL* | rs11570891 | Hypercholesterolemia | T | C | 0.95 (0.92-0.97) | 1.0x10<sup>-4</sup> | TAG(52:3;0) - | *LPL* + (nerve-tibial) |
| *LPL* | rs11570891 | Ischemic heart diseases | T | C | 0.95 (0.92-0.98) | 1.1x10<sup>-4</sup> | TAG(52:3;0) - | *LPL* + (nerve-tibial) |
| *LPL* | rs11570891 | Myocardial infarction | T | C | 0.93 (0.89-0.97) | 5.3x10<sup>-4</sup> | TAG(52:3;0) - | *LPL* + (nerve-tibial) |
| *FADS2* | rs28456 | Cholelithiasis | G | A | 1.08 (1.05-1.10) | 5.7x10<sup>-9</sup> | CE(20:4;0) - | *FADS2* + (blood)<br>*FADS1* - (pancreas) |
| *FADS2* | rs28456 | Phlebitis and thrombophlebitis | G | A | 0.92 (0.88-0.96) | 3.9x10<sup>-4</sup> | CE(20:4;0) - | *FADS2* + (blood)<br>*FADS1* - (pancreas) |
| *APOA5* | rs964184 | Hypercholesterolemia | C | G | 0.86 (0.84-0.89) | 1.5x10<sup>-32</sup> | TAG(52:3;0) - |  |
| *APOA5* | rs964184 | Hypertension | C | G | 0.95 (0.93-0.97) | 7.2x10<sup>-8</sup> | TAG(52:3;0) - |  |
| *APOA5* | rs964184 | Coronary atherosclerosis | C | G | 0.94 (0.91-0.97) | 1.3x10<sup>-4</sup> | TAG(52:3;0) - |  |
| *SPTLC3* | rs364585 | Cholelithiasis | G | A | 0.95 (0.93-0.98) | 1.2x10<sup>-4</sup> | Total CER - | *SPTLC3* - (liver) |

The PC(16:0;0-16:0;0) associated locus-*BLK* (rs1478898-A), which is an eQTL for *BLK*, showed association with decreased risk of obesity and hypertension, and increased risk of gallbladder disorders. In addition to its role in B-cell receptor signaling and B-cell development, *BLK* stimulates insulin synthesis and secretion in response to glucose and enhances the expression of several pancreatic beta-cell transcription factors.^37^ Consistent to its physiological roles, *BLK* has previously been implicated in autoimmune diseases such as systemic lupus erythematosus,^38^ maturity-onset diabetes of the young (MODY),^37^ and hypertension,^39^ however its role in obesity and gallbladder disorder has not been described before.

*SPTLC3* rs364585-G (associated with reduced levels of ceramides) showed association with reduced risk of gallbladder disorders (cholelithiasis). *SPTLC3* is a rate-limiting enzyme involved in ceramide biosynthesis. Inhibition of ceramides biosynthesis has been suggested to suppress the gallstones formation in animal model,^40^ however the relationship has not been explored in humans yet. Our results suggest that rs364585-G, which is an eQTL for *SPTLC3* leading to reduced expression of *SPTLC3*, might modulate the risk of gallbladder disorders through its effect on ceramide biosynthesis. Altogether, our PheWAS results suggest potential disease susceptibility genes that warrant further investigation to validate and understand their role in disease outcomes.

### Lipidomics provide higher statistical power

As intermediate phenotypes are known to provide more statistical power, we assessed whether the lipid species could help to detect genetic associations with greater power than traditional lipid measures using variants previously identified for traditional lipid measures (number of variants=557). We found that molecular lipid species have much stronger associations than traditional lipid measures with the same sample size, except for well-known *APOE* and *CETP* (Figure 6, Supplementary Table 10). The associations were several orders of magnitudes stronger for the variants in or near genes involved in lipid metabolism such as *FADS1-2-3*, *LIPC*, *ABCG5/8*, *SGPP1*, *SPTLC3*. This shows that the lipidomics provides higher chances to identify lipid-modulating variants, particularly the ones with direct role in lipid metabolism, with much smaller sample size than traditional lipid measures.

**Figure 6:**
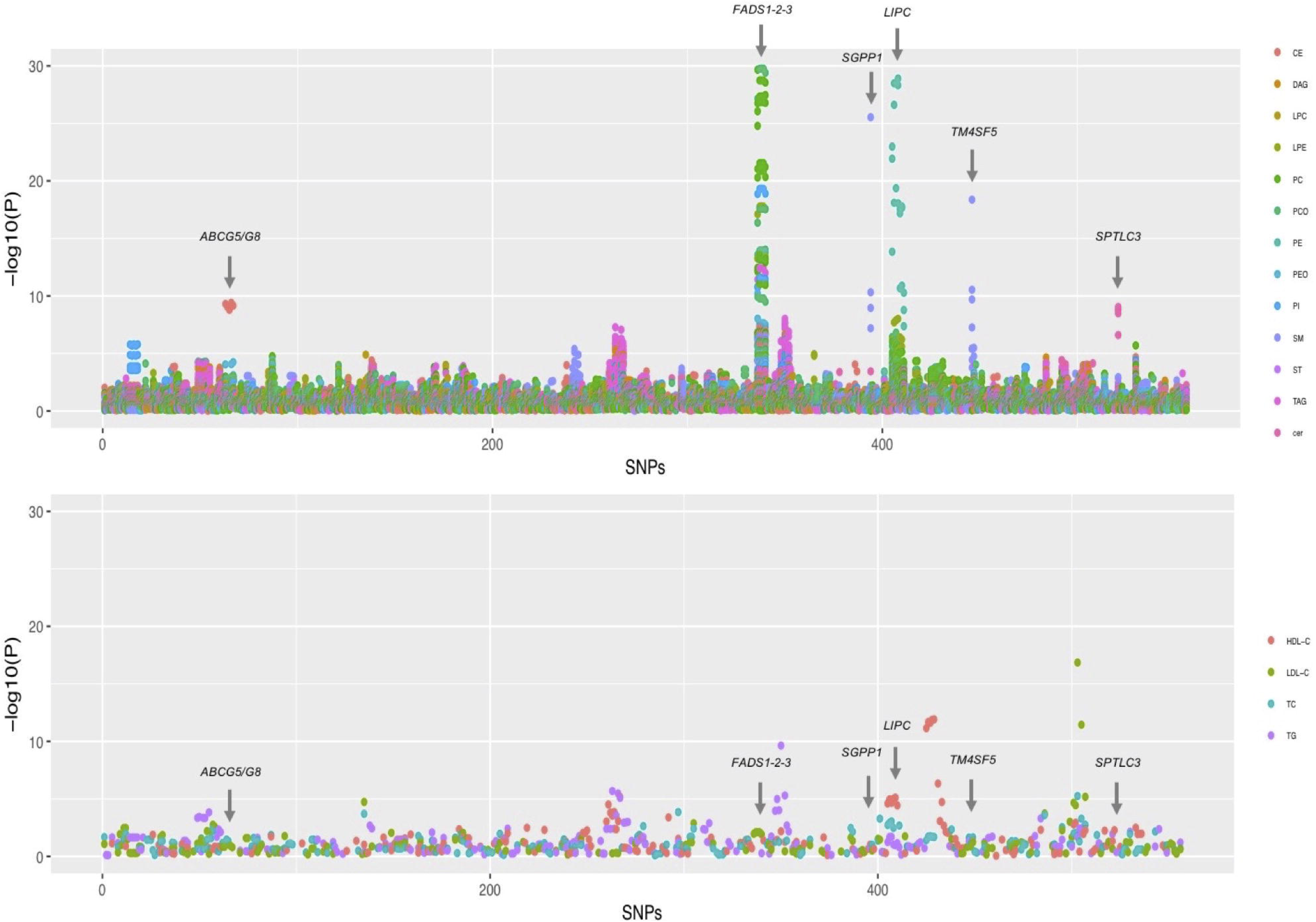
Association of known variants for traditional lipids with lipid species and traditional lipid measures. The P values for the associations of the lead SNPs (557 SNPs available in our dataset) identified through different genome-wide or exome-wide studies of traditional lipid measures (HDL-C, LDL-C, TG, TC) with lipid species (upper panel) and traditional lipid measures (lower panel) are plotted. The SNPs on the x-axis are serially arranged based on their chromosomal positions and as listed in the Supplementary Table 9. The points on the plots are color coded by the lipid classes in upper panel and traditional lipid measure in lower panel.

## Discussion

Our study integrates lipidome, genome and phenome to reveal detailed description of genetic regulation of lipid metabolism and its effect on disease risk. To the best of our knowledge, this is the first large-scale study of genetics of lipidomics presenting the SNP based heritability, genetic sharing of the lipid species, and new genomic loci associated with one or several lipid species and disease risks in humans. The detailed profiling also provided clues to probable molecular mechanisms for genetic variants both at new and previously reported loci.

The results presented here allow us to draw several conclusions. First, despite the influence of dietary intake on the circulatory levels of lipids, plasma levels of lipid species are found to be heritable, suggesting considerable role of endogenous regulation in lipid metabolism. Importantly, genetic mechanisms seem not to regulate all lipid species in a lipid class in the same way, as also observed in recent mice lipidomics studies.^41,42^ Longer and more unsaturated lipid species from different lipid classes clearly display a greater genetic sharing. These observations are consistent with a previous study based on family pedigree.^33^ Our finding is important in the light of the proposed role of lipids containing PUFAs in CVD, diabetes, and neurological disorders.^43–45^ Identification of genetic factors regulating these particular lipids is important for understanding the subtleties of lipid metabolism and devising disease preventive strategies; including dietary interventions for these common complex diseases that cause an enormous public health burden globally. Our study provides multiple leads in this direction by identifying 11 genomic loci (*KLHL17*, *APOA1*, *CD33*, *SHTN1*, *FADS2*, *LIPC*, *MBOAT7, MIR100HG*, *PTPRN1*, and *TMEM86B*) associated with long, polyunsaturated lipids. Of these, *FADS2*, *APOA1*, and *LPL* variants were also associated with cardiovascular related phenotypes in our PheWAS analysis (Figure 5). We discuss below how identification of genetic variants in these lipid species associated loci has helped us to provide new insights to their role in lipid metabolism.

Second, we identified genetic variants associated with 74 lipid species from 11 lipid classes. Individual lipid species from several lipid classes including CERs, CEs, TAGs, and PCs have been shown to predict risk for CVD and diabetes.^4–10^ This knowledge can directly fuel studies on disease markers or drug target discovery. For example, Cer(d18:1/24:0) is recently reported to be associated with increased risk of CVD events.^46^ We have identified a variant associated with Cer(42:1;2) (this species presumably includes Cer(d18:1/24:0) molecular species) near *ZNF385D*. The *ZNF385D* rs13070110-C was associated with increased levels of Cer(42:1;2). We also observed nominal association of rs13070110-C with increased risk of arterial and venous thrombosis (Supplementary Table 9). CEs have also been reported to modulate the risk of CVD events.^4,8^ Our study revealed three loci associated with CEs, including two novel loci-*ABCG5/8* and *SYNGR1*. The rs76866386-C at *ABCG5/8*, which codes for ABC sterol transporters G5 and G8, has previously been associated with TC, LDL-C and CEs in LDL,^11,16^ however we identified a specific association of rs76866386C with reduced level of cholesteryl ester species CE(20:2;0). CE(20:2;0) has been shown to be positively associated with high LDL-C,^47^ suggesting that the previously observed associations of rs76866386 with LDL-C and CE in LDL may have been contributed by the specific lipid species CE(20:2;0).

Third, our genetic associations give several clues to new biology, both in new and known lipid loci. With a few examples, we show here how detailed profiling of molecular lipid species can help us understand the effects of genetic variants or genes in specific metabolic pathways at molecular level. Genetic variants at *FADS* gene cluster have been consistently reported to be associated with omega-3 and omega-6 fatty acids levels with inverse effects on different PUFAs, however, its mechanism has not been fully deciphered. Linoleic acid and alpha-linolenic acid are essential fatty acids and substrates for delta-6 desaturation by FADS2.^48^ This desaturation step generates gamma-linolenic acid and stearidonic acid that by elongation yields dihomo-gamma-linolenic acid and eicosatetraenoic acid (Figure 5). Further, delta-5 desaturation of dihomo-gamma-linolenic acid by *FADS1* generates arachidonic acid and eicosapentaenoic acid. Subsequent elongations and desaturations generate docosatetraenoic acid, docosapentaenoic acid, and docosahexaenoic acid. Thus, as depicted in Figure 5, the inverse efffects of *FADS2* rs28456-G on *FADS2* and *FADS1* expressions explain its effects on different PUFAs with opposite directions.

Our integrated approach provided first clue to the probable variable efficiency of LPL and APOA5 in lipolysis of TAGs. *LPL* codes for lipoprotein lipase that is the master lipolytic factor of TAGs in TAG-enriched chylomicrons and VLDL particles, whereas *APOA5* codes for the activator that stimulates LPL-mediated lipolysis of TAG-rich lipoproteins and their remnants. Our results suggest that LPL and APOA5 might have varied effects on different TAG species with more efficient lipolysis of medium length TAGs (Figure 5). It is interesting that both *LPL* and *APOA5* genetic variants had very similar association patterns with hypercholesterolemia and ischemic heart diseases also.

Similarly, lipidomic profiles helped to understand the physiological roles of variants in the newly identified genes. The patterns of associations of *PTPRN2* rs10281741-G and *MIR100HG* rs10790495-G with reduced levels of long, polyunsaturated TAG species, suggested their probable role in lipid metabolism. *PTPRN2* codes for protein tyrosine phosphatase receptor N2 with a possible role in pancreatic insulin secretion and development of diabetes mellitus.^49^ *MIR100HG* codes for long non-coding RNAs that act as regulators of cell proliferation.^50^ The *MIR100HG* rs10790495 is an eQTL for the heat shock protein HSPA8 that also has a role in cell proliferation.^51^ However, it is not known if *PTPRN2* and *MIR100HG* or *HSPA8* have any role in lipid metabolism. Our results suggest that these variants might have role in negative regulation of either elongation and desaturation of fatty acids or incorporation of long chain unsaturated fatty acids during TAG biosynthesis.

Fourth, our results point to probable risk/protective variants for diseases, highlighting the potential of using detailed lipidomics profiles in disease gene mapping. Two lipid species associated loci-*BLK* and *SPTLC3* were found to be associated with cholelithiasis (gallstones). *BLK* was also identified as a new susceptibility locus for obesity in the present study. Cholelithiasis is one of the most prevalent gastrointestinal diseases with up to 15% prevalence in adult populations. Although up to two thirds of patients do not suffer any symptoms, cholelithiasis is the most significant risk factor for acute cholecystitis. ^52^ Risk factors of cholelithiasis include obesity, hyperlipidemia and type 2 diabetes. Also, the pathogenesis of cholelithiasis is now recognized to be influenced by the immune system.^53^ Owing to its role in immune response, insulin synthesis and insulin secretion, *BLK* seems to be a potential risk modifying gene for obesity and cholelithiasis. Relationship between ceramides and cholelithiasis has also been suggested previously,^40^ and given the role of *SPTLC3* in ceramide biosynthesis, the *SPTLC3* variant might influence the risk for gallstones. However, the associations of these variants with disease risks warrant further investigation.

Finally, we show that genetic investigation utilizing only those lipid measures that are traditionally used in routine clinical work would fail to capture the association with molecular lipid species which are potential independent disease risk factors. For example, LPCs and PCs have previously been associated with incident coronary heart disease risk.^4–6^ But we show that these lipid measures have very low phenotypic and genotypic correlations with traditional lipid measures and known variants for traditional lipid measures explain only a small percentage of variance in their plasma levels. This again highlights the potential of lipidomic profiles in disease gene identification. We also demonstrate that individual lipid species have stronger statistical power to detect the associations compared to the traditional lipid measures at all known lead variants using the same dataset. These findings suggest that lipidomic profiles capture information beyond standard lipid measures and provide opportunity to identify additional genetic variants influencing lipid metabolism.

Our study had some potential limitations. To the best of our knowledge, the present study is the largest genetic screen of lipidomic variation, but larger cohorts are needed to achieve full understanding of the genetic regulation of detailed lipid metabolism. The EUFAM cohort samples were from fasting state whereas the FINRISK cohort samples were semi-fasting. This does not seem to have substantial effect on lipidomic profiles as we demonstrated similar lipidomic profiles for dyslipidemias from the EUFAM and FINRISK cohorts.^47^ Moreover, the recent guidelines from the European Atherosclerosis Society and European Federation of Clinical Chemistry and Laboratory Medicine recommends non-fasting blood samples for assessment of plasma lipid profiles.^54^ The UK Biobank cohort is reported to have “healthy volunteer” effect,^55^ which may affect the PheWAS results. However, given the large sample size, the selection bias is unlikely to have substantial effect on genetic case-control association analyses. Furthermore, lipidomic profiles were measured in whole plasma comprising of all lipoprotein classes and particle sizes, which does not provide information at the level of individual lipoprotein subclasses and limits our ability to gain detailed mechanistic insights. We also excluded poorly detected lipid species from all the analyses to ensure high data quality that narrowed the spectrum of lipidomic profiles. Further advances in lipidomics platforms might help to capture more comprehensive and complete lipidomic profiles, including the position of fatty acyl chains in glycerol backbone of TAGs and glycerophospholipids and detection of sphingosine-1-P species and several other species, that would allow to overcome these limitations.

In conclusion, our study demonstrates that lipidomics enable deeper insights to the genetic regulation of lipid metabolism than clinically used lipid measures, which in turn might help guide future biomarker and drug target discovery and disease prevention.

## Funding

This work was supported by National Institutes of Health [grant HL113315 to SR, MRT and AP]; Finnish Foundation for Cardiovascular Research [to SR, VS, MRT, MJ, and AP]; Academy of Finland Center of Excellence in Complex Disease Genetics [grant 312062 to SR]; Academy of Finland [285380 to SR, 288509 to MP]; Jane and Aatos Erkko Foundation [to MJ]; Sigrid Jusélius Foundation [to SR and MRT]; Horizon 2020 Research and Innovation Programme [grant 692145 to SR]; EU-project RESOLVE (EU 7th Framework Program) [grant 305707 to MRT]; HiLIFE Fellowship [to SR]; Helsinki University Central Hospital Research Funds [to MRT]; Magnus Ehrnrooth Foundation [to MJ]; Leducq Foundation [to MRT]; Ida Montin Foundation [to PR]; MD-PhD Programme of the Faculty of Medicine, University of Helsinki [to JTR]; Doctoral Programme in Population Health, University of Helsinki [to JTR and PR]; Finnish Medical Foundation [to JTR]; Emil Aaltonen Foundation [to JTR and PR]; Biomedicum Helsinki Foundation [to JTR]; Paulo Foundation [to JTR]; Idman Foundation [to JTR]; Veritas Foundation [to JTR]; FIMMEMBL PhD Fellowship grant [to SH]. The funders had no role in study design, data collection and analysis, decision to publish, or preparation of the manuscript.

## Acknowledgements

We would like to thank Sari Kivikko, Huei-Yi Shen, and Ulla Tuomainen for management assistance. We thank all study participants for their participation. The FINRISK data used for the research were obtained from THL Biobank. We thank the THL DNA laboratory for its skillful work to produce the DNA samples used in the genotyping work, which was used in this study. Part of the genotyping was performed by the Institute for Molecular Medicine Finland FIMM Technology Centre, University of Helsinki. This research has been conducted using the UK Biobank Resource with application number 22627.

## Conflict of Interest

VS has participated in a conference trip sponsored by Novo Nordisk and received an honorarium from the same source for participating in an advisory board meeting. MJG is employee of Lipotype GmbH, CK is shareholder and employee of Lipotype GmbH, KS is shareholder and CEO of Lipotype GmbH. MAS is a shareholder of Lipotype GmbH and an employee of PORT.

## References

1. Ference BA, Ginsberg HN, Graham I, Ray KK, Packard CJ, Bruckert E, Hegele RA, Krauss RM, Raal FJ, Schunkert H, Watts GF, Boren J, Fazio S, Horton JD, Masana L, Nicholls SJ, Nordestgaard BG, van de Sluis B, Taskinen MR, Tokgöoglu L, Landmesser U, Laufs U, Wiklund O, Stock JK, Chapman MJ, Catapano AL. Low-density lipoproteins cause atherosclerotic cardiovascular disease. 1. Evidence from genetic, epidemiologic, and clinical studies. A consensus statement from the european atherosclerosis society consensus panel. Eur Heart J. 2017;38:2459–2472.

2. Schulze MB, Weikert C, Pischon T, Bergmann MM, Al-Hasani H, Schleicher E, Fritsche A, Häring HU, Boeing H, Joost HG. Use of multiple metabolic and genetic markers to improve the prediction of type 2 diabetes: the EPIC-Potsdam Study. Diabetes Care. 2009;32:2116–2119.

3. Quehenberger O, Dennis EA. The human plasma lipidome. N Engl J Med. 2011;365:1812–1823.

4. Stegemann C, Pechlaner R, Willeit P, Langley SR, Mangino M, Mayr U, Menni C, Moayyeri A, Santer P, Rungger G, Spector TD, Willeit J, Kiechl S, Mayr M. Lipidomics profiling and risk of cardiovascular disease in the prospective population-based bruneck study. Circulation. 2014;129:1821–1831.

5. Alshehry ZH, Mundra PA, Barlow CK, Mellett NA, Wong G, McConville MJ, Simes J, Tonkin AM, Sullivan DR, Barnes EH, Nestel PJ, Kingwell BA, Marre M, Neal B, Poulter NR, Rodgers A, Williams B, Zoungas S, Hillis GS, Chalmers J, Woodward M, Meikle PJ. Plasma lipidomic profiles improve on traditional risk factors for the prediction of cardiovascular events in type 2 diabetes mellitus. Circulation. 2016;134:1637–1650.

6. Laaksonen R, Ekroos K, Sysi-Aho M, Hilvo M, Vihervaara T, Kauhanen D, Suoniemi M, Hurme R, Marz W, Scharnagl H, Stojakovic T, Vlachopoulou E, Lokki ML, Nieminen MS, Klingenberg R, Matter CM, Hornemann T, Jüni P, Rodondi N, Räber L, Windecker S, Gencer B, Pedersen ER, Tell GS, Nygård O, Mach F, Sinisalo J, Lüscher TF. Plasma ceramides predict cardiovascular death in patients with stable coronary artery disease and acute coronary syndromes beyond ldl-cholesterol. Eur Heart J. 2016;37:1967–1976.

7. Havulinna AS, Sysi-Aho M, Hilvo M, Kauhanen D, Hurme R, Ekroos K, Salomaa V, Laaksonen R. Circulating Ceramides Predict Cardiovascular Outcomes in the Population-Based FINRISK 2002 Cohort. Arterioscler Thromb Vasc Biol. 2016;36:2424–2430.

8. Razquin C, Liang L, Toledo E, Clish CB, Ruiz-Canela M, Zheng Y, Wang DD, Corella D, Castaner O, Ros E, Aros F, Gomez-Gracia E, Fiol M, Santos-Lozano JM, Guasch-Ferre M, Serra-Majem L, Sala-Vila A, Buil-Cosiales P, Bullo M, Fito M, Portoles O, Estruch R, Salas Salvado J, Hu FB, Martinez-Gonzalez MA. Plasma lipidome patterns associated with cardiovascular risk in the PREDIMED trial: A case-cohort study. Int J Cardiol. 2018;253:126–132.

9. Hilvo M, Salonurmi T, Havulinna AS, Kauhanen D, Pedersen ER, Tell GS, Meyer K, Teeriniemi AM, Laatikainen T, Jousilahti P, Savolainen MJ, Nygård O, Salomaa V, Laaksonen R. Ceramide stearic to palmitic acid ratio predicts incident diabetes. Diabetologia. 2018;61:1424–1434.

10. Khaw KT, Friesen MD, Riboli E, Luben R, Wareham N. Plasma phospholipid fatty acid concentration and incident coronary heart disease in men and women: the EPIC-Norfolk prospective study. PLoS Med. 2012;9:e1001255.

11. Surakka I, Horikoshi M, Mägi R, Sarin AP, Mahajan A, Lagou V. The impact of low-frequency and rare variants on lipid levels. Nat Genet. 2015;47:589–97.

12. Liu DJ, Peloso GM, Yu H, Butterworth AS, Wang X, Mahajan A, Saleheen D, Emdin C, Alam D, Alves AC, Amouyel P, Di Angelantonio E, Arveiler D, Assimes TL, Auer PL, Baber U, Ballantyne CM, Bang LE, Benn M, Bis JC, Boehnke M, Boerwinkle E, Bork-Jensen J, Bottinger EP, Brandslund I, Brown M, Busonero F, Caulfield MJ, Chambers JC, Chasman DI, Chen YE, Chen YI, Chowdhury R, Christensen C, Chu AY, Connell JM, Cucca F, Cupples LA, Damrauer SM, Davies G, Deary IJ, Dedoussis G, Denny JC, Dominiczak A, Dubé MP, Ebeling T, Eiriksdottir G, Esko T, Farmaki AE, Feitosa MF, Ferrario M, Ferrieres J, Ford I, Fornage M, Franks PW, Frayling TM, Frikke-Schmidt R, Fritsche LG, Frossard P, Fuster V, Ganesh SK, Gao W, Garcia ME, Gieger C, Giulianini F, Goodarzi MO, Grallert H, Grarup N, Groop L, Grove ML, Gudnason V, Hansen T, Harris TB, Hayward C, Hirschhorn JN, Holmen OL, Huffman J, Huo Y, Hveem K, Jabeen S, Jackson AU, Jakobsdottir J, Jarvelin MR, Jensen GB, Jørgensen ME, Jukema JW, Justesen JM, Kamstrup PR, Kanoni S, Karpe F, Kee F, Khera AV, Klarin D, Koistinen HA, Kooner JS, Kooperberg C, Kuulasmaa K, Kuusisto J, Laakso M, Lakka T, Langenberg C, Langsted A, Launer LJ, Lauritzen T, Liewald DCM, Lin LA, Linneberg A, Loos RJF, Lu Y, Lu X, Mägi R, Malarstig A, Manichaikul A, Manning AK, Mäntyselkä P, Marouli E, Masca NGD, Maschio A, Meigs JB, Melander O, Metspalu A, Morris AP, Morrison AC, Mulas A, Müller-Nurasyid M, Munroe PB, Neville MJ, Nielsen JB, Nielsen SF, Nordestgaard BG, Ordovas JM, Mehran R, O’Donnell CJ, Orho-Melander M, Molony CM, Muntendam P, Padmanabhan S, Palmer CNA, Pasko D, Patel AP, Pedersen O, Perola M, Peters A, Pisinger C, Pistis G, Polasek O, Poulter N, Psaty BM, Rader DJ, Rasheed A, Rauramaa R, Reilly DF, Reiner AP, Renström F, Rich SS, Ridker PM, Rioux JD, Robertson NR, Roden DM, Rotter JI, Rudan I, Salomaa V, Samani NJ, Sanna S, Sattar N, Schmidt EM, Scott RA, Sever P, Sevilla RS, Shaffer CM, Sim X, Sivapalaratnam S, Small KS, Smith AV, Smith BH, Somayajula S, Southam L, Spector TD, Speliotes EK, Starr JM, Stirrups KE, Stitziel N, Strauch K, Stringham HM, Surendran P, Tada H, Tall AR, Tang H, Tardif JC, Taylor KD, Trompet S, Tsao PS, Tuomilehto J, Tybjaerg-Hansen A, van Zuydam NR, Varbo A, Varga TV, Virtamo J, Waldenberger M, Wang N, Wareham NJ, Warren HR, Weeke PE, Weinstock J, Wessel J, Wilson JG, Wilson PWF, Xu M, Yaghootkar H, Young R, Zeggini E, Zhang H, Zheng NS, Zhang W, Zhang Y, Zhou W, Zhou Y, Zoledziewska M; Charge Diabetes Working Group; EPIC-InterAct Consortium; EPIC-CVD Consortium; GOLD Consortium; VA Million Veteran Program, Howson JMM, Danesh J, McCarthy MI, Cowan CA, Abecasis G, Deloukas P, Musunuru K, Willer CJ, Kathiresan S. Exome-wide association study of plasma lipids in >300,000 individuals. Nat Genet. 2017;49:1758–1766.

13. Demirkan A, van Duijn CM, Ugocsai P, Isaacs A, Pramstaller PP, Liebisch G, Wilson JF, Johansson Å, Rudan I, Aulchenko YS, Kirichenko AV, Janssens AC, Jansen RC, Gnewuch C, Domingues FS, Pattaro C, Wild SH, Jonasson I, Polasek O, Zorkoltseva IV, Hofman A, Karssen LC, Struchalin M, Floyd J, Igl W, Biloglav Z, Broer L, Pfeufer A, Pichler I, Campbell S, Zaboli G, Kolcic I, Rivadeneira F, Huffman J, Hastie ND, Uitterlinden A, Franke L, Franklin CS, Vitart V; Diagram Consortium, Nelson CP, Preuss M; Cardiogram Consortium, Bis JC, O’Donnell CJ, Franceschini N; Charge Consortium, Witteman JC, Axenovich T, Oostra BA, Meitinger T, Hicks AA, Hayward C, Wright AF, Gyllensten U, Campbell H, Schmitz G; EUROSPAN consortium. Genome-wide association study identifies novel loci associated with circulating phospho-and sphingolipid concentrations. PLoS Genet. 2012;8:e1002490.

14. Hicks AA, Pramstaller PP, Johansson A, Vitart V, Rudan I, Ugocsai P, Aulchenko Y, Franklin CS, Liebisch G, Erdmann J, Jonasson I, Zorkoltseva IV, Pattaro C, Hayward C, Isaacs A, Hengstenberg C, Campbell S, Gnewuch C, Janssens AC, Kirichenko AV, König IR, Marroni F, Polasek O, Demirkan A, Kolcic I, Schwienbacher C, Igl W, Biloglav Z, Witteman JC, Pichler I, Zaboli G, Axenovich TI, Peters A, Schreiber S, Wichmann HE, Schunkert H, Hastie N, Oostra BA, Wild SH, Meitinger T, Gyllensten U, van Duijn CM, Wilson JF, Wright A, Schmitz G, Campbell H. Genetic Determinants of Circulating Sphingolipid Concentrations in European Populations. PLoS Genet. 2009;5:e1000672.

15. Gieger C, Geistlinger L, Altmaier E, Hrabé de Angelis M, Kronenberg F, Meitinger T, Mewes HW, Wichmann HE, Weinberger KM, Adamski J, Illig T, Suhre K. Genetics meets metabolomics: a genome-wide association study of metabolite profiles in human serum. PLoS Genet. 2008;4:e1000282.

16. Kettunen J, Demirkan A, Würtz P, Draisma HH, Haller T, Rawal R, Vaarhorst A, Kangas AJ, Lyytikäinen LP, Pirinen M, Pool R, Sarin AP, Soininen P, Tukiainen T, Wang Q, Tiainen M, Tynkkynen T, Amin N, Zeller T, Beekman M, Deelen J, van Dijk KW, Esko T, Hottenga JJ, van Leeuwen EM, Lehtimäki T, Mihailov E, Rose RJ, de Craen AJ, Gieger C, Kähönen M, Perola M, Blankenberg S, Savolainen MJ, Verhoeven A, Viikari J, Willemsen G, Boomsma DI, van Duijn CM, Eriksson J, Jula A, Järvelin MR, Kaprio J, Metspalu A, Raitakari O, Salomaa V, Slagboom PE, Waldenberger M, Ripatti S, Ala-Korpela M. Genome-wide study for circulating metabolites identifies 62 loci and reveals novel systemic effects of LPA. Nat Commun. 2016;7:11122.

17. Rhee EP, Ho JE, Chen MH, Shen D, Cheng S, Larson MG, Ghorbani A, Shi X, Helenius IT, O’Donnell CJ, Souza AL, Deik A, Pierce KA, Bullock K, Walford GA, Vasan RS, Florez JC, Clish C, Yeh JR, Wang TJ, Gerszten RE. A genome-wide association study of the human metabolome in a community-based cohort. Cell Metab. 2013;18:130–143.

18. Draisma HHM, Pool R, Kobl M, Jansen R, Petersen AK, Vaarhorst AAM, Yet I, Haller T, Demirkan A, Esko T, Zhu G, Böhringer S, Beekman M, van Klinken JB, Römisch-Margl W, Prehn C, Adamski J, de Craen AJM, van Leeuwen EM, Amin N, Dharuri H, Westra HJ, Franke L, de Geus EJC, Hottenga JJ, Willemsen G, Henders AK, Montgomery GW, Nyholt DR, Whitfield JB, Penninx BW, Spector TD, Metspalu A, Slagboom PE, van Dijk KW, ‘t Hoen PAC, Strauch K, Martin NG, van Ommen GB, Illig T, Bell JT, Mangino M, Suhre K, McCarthy MI, Gieger C, Isaacs A, van Duijn CM, Boomsma DI. Genome-wide association study identifies novel genetic variants contributing to variation in blood metabolite levels. Nat Commun. 2015;6:7208.

19. Illig T, Gieger C, Zhai G, Römisch-Margl W, Wang-Sattler R, Prehn C, Altmaier E, Kastenmüller G, Kato BS, Mewes HW, Meitinger T, de Angelis MH, Kronenberg F, Soranzo N, Wichmann HE, Spector TD, Adamski J, Suhre K. A genome-wide perspective of genetic variation in human metabolism. Nat Genet. 2010;42:137–141.

20. Shin SY, Fauman EB, Petersen AK, Krumsiek J, Santos R, Huang J, Arnold M, Erte I, Forgetta V, Yang TP, Walter K, Menni C, Chen L, Vasquez L, Valdes AM, Hyde CL, Wang V, Ziemek D, Roberts P, Xi L, Grundberg E; Multiple Tissue Human Expression Resource (MuTHER) Consortium, Waldenberger M, Richards JB, Mohney RP, Milburn MV, John SL, Trimmer J, Theis FJ, Overington JP, Suhre K, Brosnan MJ, Gieger C, Kastenmüller G, Spector TD, Soranzo N. An atlas of genetic influences on human blood metabolites. Nat Genet. 2014;46:543–550.

21. Porkka KV, Nuotio I, Pajukanta P, Ehnholm C, Suurinkeroinen L, Syvänne M, Lehtimäki T, Lahdenkari AT, Lahdenperä S, Ylitalo K, Antikainen M, Perola M, Raitakari OT, Kovanen P, Viikari JS, Peltonen L, Taskinen MR. Phenotype expression in familial combined hyperlipidemia. Atherosclerosis. 1997;133:245–253.

22. Borodulin K, Vartiainen E, Peltonen M, Jousilahti P, Juolevi A, Laatikainen T, Männistö S, Salomaa V, Sundvall J, Puska P. Forty-year trends in cardiovascular risk factors in Finland. Eur J Public Health. 2015;25:539–546.

23. Sudlow C, Gallacher J, Allen N, Beral V, Burton P, Danesh J, Downey P, Elliott P, Green J, Landray M, Liu B, Matthews P, Ong G, Pell J, Silman A, Young A, Sprosen T, Peakman T, Collins R. UK Biobank: An Open Access Resource for Identifying the Causes of a Wide Range of Complex Diseases of Middle and Old Age. PLoS Med. 2015;12:e1001779.

24. B. Howie, C. Fuchsberger, M. Stephens, J. Marchini, and G. R. Abecasis. Fast and accurate genotype imputation in genome-wide association studies through pre-phasing. Nat Genet. 2012;44:955–959.

25. Bycroft C, Freeman C, Petkova D, Band G, Elliott LT, Sharp K, Motyer, A, Vukcevic D, Delaneau O, O’Connell J, Cortes A, Welsh S, McVean G, Leslie S, Donnelly P, Marchini J. Genome-wide genetic data on ~500,000 UK Biobank participants. bioRxiv, 2017. 166298. https://doi.org/10.1101/166298

26. Zaitlen N, Kraft P, Patterson N, Pasaniuc B, Bhatia G, Pollack S, Price AL. Using extended genealogy to estimate components of heritability for 23 quantitative and dichotomous traits. PLoS Genet. 2013;9:e1003520.

27. Pirinen M, Benner C, Marttinen P, Järvelin MR, Rivas MA, Ripatti S. biMM: efficient estimation of genetic variances and covariances for cohorts with high-dimensional phenotype measurements. Bioinformatics. 2017;33:2405–2407.

28. Pirinen M, Donnelly P, Spencer CCA. Efficient Computation with a Linear Mixed Model on Large-scale Data Sets with Applications to Genetic Studies. Ann Appl Stat. 2012;7:369–390.

29. Marchini J., Howie B. Genotype imputation for genome-wide association studies. Nat Rev Genet. 2010;11:499–511.

30. Willer CJ, Li Y, Abecasis GR. METAL: fast and efficient meta-analysis of genomewide association scans. Bioinformatics. 2010;26:2190–2191.

31. Dey R, Schmidt EM, Abecasis GR, Lee S. A Fast and Accurate Algorithm to Test for Binary Phenotypes and Its Application to PheWAS. Am J Hum Genet. 2017;101:37–49.

32. Zhou W, Nielsen JB, Fritsche LG, Dey R, Gabrielsen ME, Wolford BN, LeFaive J, VandeHaar P, Gagliano SA, Gifford A, Bastarache LA, Wei WQ, Denny JC, Lin M,Hveem K, Kang HM, Abecasis GR, Willer CJ, Lee S. Efficiently controlling for case-control imbalance and sample relatedness in large-scale genetic association studies. Nat Genet. 2018;50:1335–1341.

33. Bellis C, Kulkarni H, Mamtani M, Kent JW Jr, Wong G, Weir JM, Barlow CK, Diego V, Almeida M, Dyer TD, Göring HHH, Almasy L, Mahaney MC, Comuzzie AG, Williams-Blangero S, Meikle PJ, Blangero J, Curran JE. Human plasma lipidome is pleiotropically associated with cardiovascular risk factors and death. Circ Cardiovasc Genet. 2014;7:854–863.

34. Frahnow T, Osterhoff MA, Hornemann S, Kruse M, Surma MA, Klose C, Simons K, Pfeiffer AFH. Heritability and responses to high fat diet of plasma lipidomics in a twin study. Sci Rep 2017;7:3750.

35. Hartmann S, Ridley AJ, Lutz S. The Function of Rho-Associated Kinases ROCK1 and ROCK2 in the Pathogenesis of Cardiovascular Disease. Frontiers in Pharmacology. 2015;6:276.

36. Chun KH, Choi KD, Lee DH, Jung Y, Henry RR, Ciaraldi TP, Kim YB. In vivo activation of ROCK1 by insulin is impaired in skeletal muscle of humans with type 2 diabetes. Am J Physiol Endocrinol Metab. 2011;300:E536–42.

37. Borowiec M, Liew CW, Thompson R, Boonyasrisawat W, Hu J, Mlynarski WM, El Khattabi I, Kim SH, Marselli L, Rich SS, Krolewski AS, Bonner-Weir S, Sharma A, Sale M, Mychaleckyj JC, Kulkarni RN, Doria A. Mutations at the BLK locus linked to maturity onset diabetes of the young and beta-cell dysfunction. Proc Natl Acad Sci USA. 2009;106:14460–14465.

38. Bentham J, Morris DL, Graham DSC, Pinder CL, Tombleson P, Behrens TW, Martín J, Fairfax BP, Knight JC, Chen L, Replogle J, Syvänen AC, Rönnblom L, Graham RR, Wither JE, Rioux JD, Alarcón-Riquelme ME, Vyse TJ. Genetic association analyses implicate aberrant regulation of innate and adaptive immunity genes in the pathogenesis of systemic lupus erythematosus. Nat Genet. 2015;47:1457–1464.

39. Ho JE, Levy D, Rose L, Johnson AD, Ridker PM, Chasman DI. Discovery and replication of novel blood pressure genetic loci in the Women’s Genome Health Study. J Hypertens. 2011;29:62–69.

40. Lee BJ, Kim JS, Kim BK, Jung SJ, Joo MK, Hong SG, Kim JS, Kim JH, Yeon JE, Park JJ, Byun KS, Bak YT, Yoo HS, Oh S. Effects of sphingolipid synthesis inhibition on cholesterol gallstone formation in C57BL/6J mice. J Gastroenterol Hepatol. 2010;25:1105–1110.

41. Jha P, McDevitt MT, Gupta R, Quiros PM, Williams EG, Gariani K, Sleiman MB, Diserens L, Jochem A, Ulbrich A, Coon JJ, Auwerx J, Pagliarini DJ. Systems Analyses Reveal Physiological Roles and Genetic Regulators of Liver Lipid Species. Cell Systems. 2018;6:722–733.e6.

42. Jha P, McDevitt MT, Halilbasic E, Williams EG, Quiros PM, Gariani K, Sleiman MB, Gupta R, Ulbrich A, Jochem A, Coon JJ, Trauner M, Pagliarini DJ, Auwerx J. Genetic Regulation of Plasma Lipid Species and Their Association with Metabolic Phenotypes. Cell Systems. 2018;6:709–721.e6.

43. Ander BP, Dupasquier CM, Prociuk MA, Pierce GN. Polyunsaturated fatty acids and their effects on cardiovascular disease. Experimental & Clinical Cardiology. 2003;8:164–172.

44. Forouhi NG, Imamura F, Sharp SJ, Koulman A, Schulze MB, Zheng J, Ye Z, Sluijs I, Guevara M, Huerta JM, Kröger J, Wang LY, Summerhill K, Griffin JL, Feskens EJ, Affret A, Amiano P, Boeing H, Dow C, Fagherazzi G, Franks PW, Gonzalez C, Kaaks R, Key TJ, Khaw KT, Kühn T, Mortensen LM, Nilsson PM, Overvad K, Pala V, Palli D, Panico S, Quirós JR, Rodriguez-Barranco M, Rolandsson O, Sacerdote C, Scalbert A, Slimani N, Spijkerman AM, Tjonneland A, Tormo MJ, Tumino R, van der A DL, van der Schouw YT, Langenberg C, Riboli E, Wareham NJ. Association of Plasma Phospholipid n-3 and n-6 Polyunsaturated Fatty Acids with Type 2 Diabetes: The EPIC-InterAct Case-Cohort Study. PLoS Med. 2016;13:e1002094.

45. Dyall SC. Long-chain omega-3 fatty acids and the brain: a review of the independent and shared effects of EPA, DPA and DHA. Front Aging Neurosci. 2015;7:52.

46. Wang DD, Toledo E, Hruby A, Rosner BA, Willett WC, Sun Q, Razquin C, Zheng Y, Ruiz-Canela M, Guasch-Ferré M, Corella D, Gómez-Gracia E, Fiol M, Estruch R, Ros E, Lapetra J, Fito M, Aros F, Serra-Majem L, Lee CH, Clish CB, Liang L, Salas-Salvadó J, Martínez-González MA, Hu FB. Plasma Ceramides, Mediterranean Diet, and Incident Cardiovascular Disease in the PREDIMED Trial (Prevención con Dieta Mediterránea). Circulation. 2017;135:2028–2040.

47. Ramo TJ, Ripatti P, Tabassum R, Soderlund S, Matikainen N, Gerl MJ, Klose C, Stitziel NO, Havulinna AS, Salomaa V, Freimer NB, Jauhiainen M, Palotie A, Taskinen MR, Simons K, Ripatti S. Coronary artery disease risk and lipidomic profiles are similar in familial and population-ascertained hyperlipidemias. bioRxiv. 2018. 321752; doi: https://doi.org/10.1101/321752

48. Saini RK, Keum YS. Omega-3 and omega-6 polyunsaturated fatty acids: Dietary sources, metabolism, and significance - A review. Life Sci. 2018;203:255–267.

49. Doi A, Shono T, Nishi M, Furuta H, Sasaki H, et al. IA-2beta, but not IA-2, is induced by ghrelin and inhibits glucose-stimulated insulin secretion. Proc Natl Acad Sci USA. 2006;103:885–890.

50. Wang S, Ke H, Zhang H, Ma Y, Ao L, Zou L, Yang Q, Zhu H, Nie J, Wu C, Jiao B. LncRNA MIR100HG promotes cell proliferation in triple-negative breast cancer through triplex formation with p27 loci. Cell Death Dis. 2018;9:805.

51. Rohde M, Daugaard M, Jensen MH, Helin K, Nylandsted J, Jäättelä M. Members of the heat-shock protein 70 family promote cancer cell growth by distinct mechanisms. Genes Dev. 2005;19:570–582.

52. Strasberg SM. Clinical practice. Acute calculous cholecystitis. N Engl J Med. 2008;358:2804–2811.

53. Maurer KJ, Carey MC, Fox JG. Roles of infection, inflammation, and the immune system in cholesterol gallstone formation. Gastroenterology. 2009;136:425–440.

54. Nordestgaard BG, Langsted A, Mora S, Kolovou G, Baum H, Bruckert E, Watts GF, Sypniewska G, Wiklund O, Boren J, Chapman MJ, Cobbaert C, Descamps OS, von Eckardstein A, Kamstrup PR, Pulkki K, Kronenberg F, Remaley AT, Rifai N, Ros E, Langlois M; European Atherosclerosis Society (EAS) and the European Federation of Clinical Chemistry and Laboratory Medicine (EFLM) Joint Consensus Initiative. Fasting is not routinely required for determination of a lipid profile: Clinical and laboratory implications including flagging at desirable concentration cutpoints-a joint consensus statement from the european atherosclerosis society and european federation of clinical chemistry and laboratory medicine. Clin. Chem. 2016;62:930–946

55. Fry A, Littlejohns TJ, Sudlow C, Doherty N, Adamska L, Sprosen T, Collins R, Allen NE. Comparison of Sociodemographic and Health-Related Characteristics of UK Biobank Participants With Those of the General Population. Am J Epidemiol. 2017;186:1026–1034.

